# Rapid long-distance migration of RPA on single stranded DNA occurs through intersegmental transfer utilizing multivalent interactions

**DOI:** 10.1101/2023.12.09.570606

**Authors:** Sushil Pangeni, Gargi Biswas, Vikas Kaushik, Sahiti Kuppa, Olivia Yang, Chang-Ting Lin, Garima Mishra, Yaakov Levy, Edwin Antony, Taekjip Ha

## Abstract

Replication Protein A (RPA) is a single stranded DNA (ssDNA) binding protein that coordinates diverse DNA metabolic processes including DNA replication, repair, and recombination. RPA is a heterotrimeric protein with six functional oligosaccharide/oligonucleotide (OB) domains and flexible linkers. Flexibility enables RPA to adopt multiple configurations and is thought to modulate its function. Here, using single molecule confocal fluorescence microscopy combined with optical tweezers and coarse-grained molecular dynamics simulations, we investigated the diffusional migration of single RPA molecules on ssDNA under tension. The diffusion coefficient *D* is the highest (20,000 nucleotides^2^/s) at 3 pN tension and in 100 mM KCl and markedly decreases when tension or salt concentration increases. We attribute the tension effect to intersegmental transfer which is hindered by DNA stretching and the salt effect to an increase in binding site size and interaction energy of RPA-ssDNA. Our integrative study allowed us to estimate the size and frequency of intersegmental transfer events that occur through transient bridging of distant sites on DNA by multiple binding sites on RPA. Interestingly, deletion of RPA trimeric core still allowed significant ssDNA binding although the reduced contact area made RPA 15-fold more mobile. Finally, we characterized the effect of RPA crowding on RPA migration. These findings reveal how the high affinity RPA-ssDNA interactions are remodeled to yield access, a key step in several DNA metabolic processes.

**Significance:** Replication Protein A (RPA) binds to the exposed single stranded DNA (ssDNA) during DNA metabolism. RPA dynamics are essential to reposition RPA on ssDNA and recruit downstream proteins at the bound site. Here in this work, we perform a detailed biophysical study on dynamics of yeast RPA on ssDNA. We show that RPA can diffuse on ssDNA and is affected by tension and salt. Our observations are best explained by the intersegmental transfer model where RPA can transiently bridge two distant DNA segments for its migration over long distances. We further dissect the contributions of the trimerization core of RPA and other adjacent RPA molecules on RPA migration. This study provides detailed experimental and computational insights into RPA dynamics on ssDNA.

## Introduction

Replication protein A was first reported as an important for *simian vacuolating virus 40* (SV40) replication in human cell extracts (1, 2), and is highly conserved in eukaryotes(3–6). RPA serves as a housekeeping single stranded DNA (ssDNA) binding protein that maintains ssDNA stability during numerous biological processes where ssDNA is transiently exposed (5, 6). RPA performs or influences many processes including protection of ssDNA from nucleases(7), resolving secondary structures like hairpins (8, 9), G-quadruplexes(10, 11), and R-Loops, (12). If unprotected by RPA, spontaneous mutations can occur to the exposed ssDNA(13, 14). RPA also marks the site of DNA damage and is required for the recruitment of ATR (ataxia-telangiectasia mutated- and Rad3-related) and activation of the ATR mediated Chk1 activation during DNA damage checkpoint(15, 16). RPA can interact with more than three dozen RPA interacting proteins that are essential for DNA metabolism and genome maintenance (16, 17).

RPA binds to ssDNA with very high affinity (dissociation constant K_D_<10^-10^ M) but must be removed or redistributed from ssDNA to make room for other DNA processing proteins. For example, RPA can help recruit helicases like HELB to ssDNA which in turn can facilitate the removal of RPA during DNA replication and recombination (18). Other mediator proteins such as Rad52 remodel select domains of RPA and gain access to the ssDNA (19). RPA is a heterotrimeric protein composed of RPA70, RPA32 and RPA14 subunits where the numbers denote the apparent molecular weight of each subunit. The subunits are composed of six oligosaccharide/oligonucleotide binding (OB) fold domains that are connected by flexible linkers of varying lengths. RPA70 houses OB-F, OB-A, OB-B and OB-C. OB-D resides in RPA32 along with a winged-helix (wh) domain. RPA14 harbors the final OB domain (OB-E) (**Figure 1A**). Four of these domains (OB-A, B, C, and D) primarily coordinate DNA binding and are termed DNA binding domains (DBDs: A, B, C and D). OB-F and the wh-domain coordinate protein-protein interactions in RPA70 and RPA32, respectively and are also termed protein-interaction domain (PIDs: PID^70N^ and PID^32C^) (20–23) (21, 24–26). RPA is held together as a constitutive heterotrimer through interactions between DBD-C (RPA70), DBD-D (RPA32), and OB-E (RPA14). While OB-E shows structural changes in the presence of ssDNA, no direct evidence for DNA binding has been observed(27). The DBDs and PIDs are connected by flexible linkers (Figure 1B), contributing to the structural flexibility of RPA (28).

**Figure 1:**
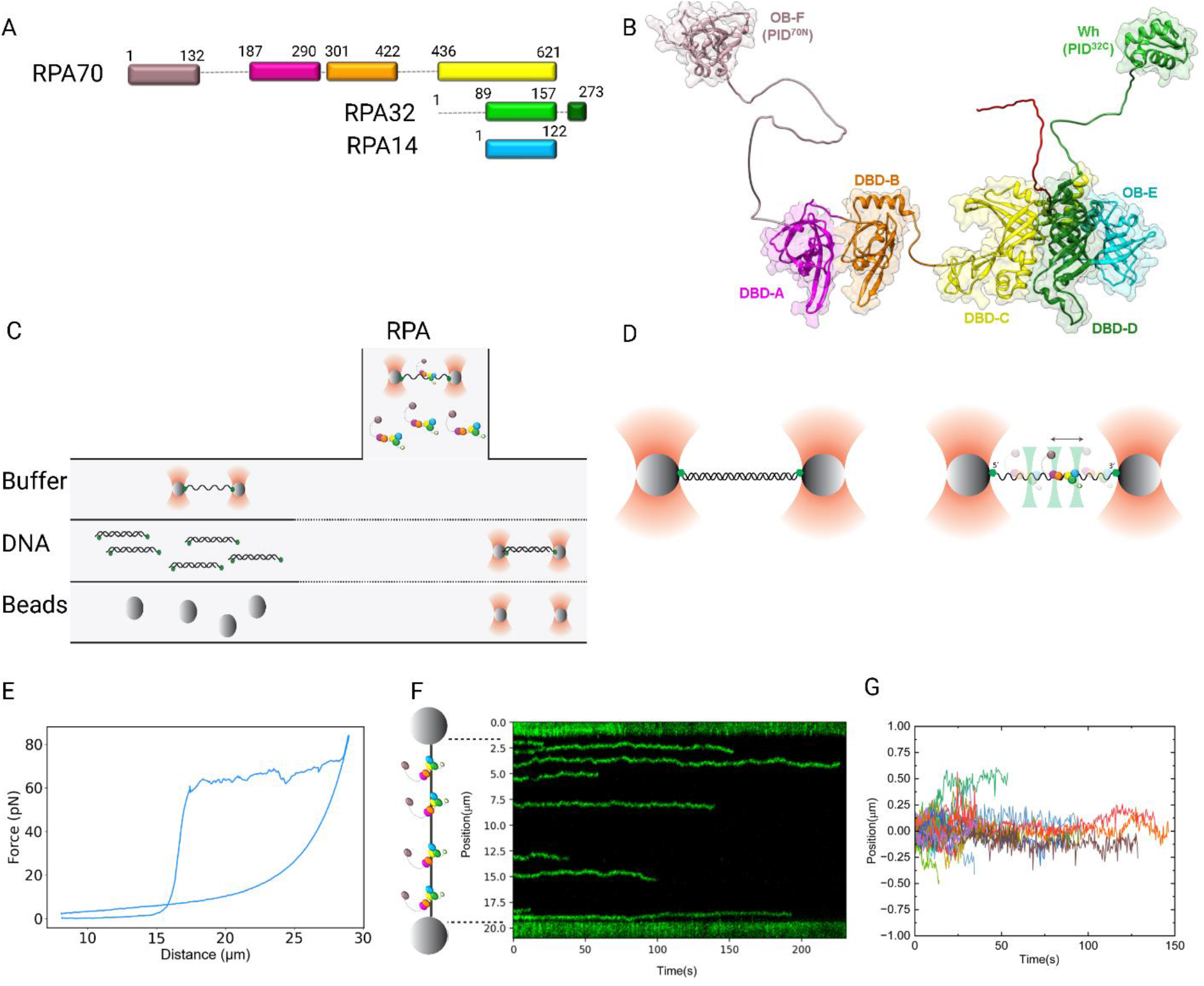
Experimental Scheme: A) Schematic of RPA subunits and their respective domains. B) A model for RPA generated from the structures of the individual OB-domains. Intervening linkers were generated using AlphaFold. ssDNA bound to the respective domains are represented as black sticks. C) Laminar flow system of commercial optical trap(C-Trap). D)Optical trap illustration. E) Force extension curve of 48.5 kbp lambda DNA to generate the ssDNA. F) Kymograph showing RPA diffusion along ssDNA. RPA DBD D diffusion along ss-DNA under 8 pN tension on ssDNA (10mM HEPES pH7.5,2mM MgCl_2,_100 mM KCl). G) Diffusion traces of many RPA molecules as in F. Diffusion start position was adjusted to position zero. Colors for each subunit of RPA match for A and B and the following illustrations.

The prevailing models for RPA-ssDNA interactions posit that the DBDs function as dynamic units with a complex array of binding, dissociation, and remodeling properties for each. Thus, RPA interacting proteins can selectively remodel one or more DBDs and PIDs and gain access to the DNA. In addition, the intrinsic flexibility of the linkers enables the DBDs and PIDs to be arranged in multiple configurations depending on the DNA substrate it encounters and the associated biological function. Recent work suggests that the overall configurational properties of RPA can be envisioned from the perspective of two halves of RPA: a dynamic and less-dynamic half(27). OB-F, DBD-A and DBD-B along with the long F-A linker (all in RPA70) are more dynamic and is positioned off the ssDNA when multiple RPA molecules bind to DNA at high density. In contrast, DBD-C (RPA70), DBD-D (RPA32), and OB-E (RPA14), which form the trimerization core (Tri-C), constitute a less-dynamic half that binds more stably to ssDNA (7, 13, 17, 27–33)(27).

An established approach to describe the DNA binding properties of RPA invokes binding mode transitions. Binding modes are defined by the occluded site size for RPA, defined as the number of nucleotides required to saturate the DNA binding site under a given condition. As each DBD can dynamically associate or dissociate from DNA, accessibility of the DNA to other interacting proteins will be dictated by the binding mode of RPA. RPA was shown to initially bind ssDNA with a low binding mode of 8-12 nt involving mainly DBD-A and DBD-B. Binding of additional DBDs (DBD-C and DBD-D) switches RPA to a higher binding mode of 17-35 nt that involves all DBDs (8, 28, 31, 34–36). A switch or transition between the binding modes can be modulated in solution by changing the ionic strength (8, 36). The low binding mode is observed at ∼ 100 mM NaCl and the higher binding mode is achieved at ∼600 mM NaCl (8). Assignment of which DBDs drive the two binding modes came from affinity measurements made with isolated DBDs. However, recent work shows that these affinity measurements are more complex when considered from the context of full-length RPA where DBDs-A and B are more dynamic (37).

Diffusion of RPA on ssDNA is another feature that has been experimentally observed (8). Yet, how RPA diffuses on long ssDNA and the underlying mechanistic basis are poorly understood. Single molecule measurements are ideally suited for highly dynamic proteins such as RPA (28, 38–44). Single molecule FRET (smFRET) (45) has been used to study RPA-ssDNA interactions and revealed the dynamic nature of RPA on ssDNA where each DBD can undergo microscopic association and dissociation while RPA is bound to the ssDNA (28, 37, 39, 40, 46) and RNA (40). Magnetic tweezers have been used to measure the force regulated dynamics of RPA at a DNA replication fork-like structure (47). Single molecule DNA curtains have also been used to study RPA dynamics (19, 38, 44, 48–50), and uncovered that RPA can undergo facilitated exchange in the presence of free excess RPA, Rad51, or SSB in solution (44, 50).

Diffusional migration of RPA on ssDNA was discovered also using single molecule methods (8, 32). Nguyen *et al* have shown diffusion of RPA along short (≤120 nt) strands of DNA using smFRET (8) and found that as the length of DNA was increased, the diffusion coefficient, *D*, can be determined more precisely (8). Although previous studies have demonstrated changes in binding mode with varying ionic strength, the effect of binding mode on the movement of RPA on DNA is unknown (8). *E. coli* single stranded DNA binding protein (SSB), which plays similar roles in bacteria, has been shown to have significantly higher diffusion coefficients on long stretches of DNA compared to short DNA, potentially due to intersegmental transfer between distant sites on DNA (51, 52). Studying RPA movement on long stretches of ssDNA will help us to examine the effect of the presence of other proteins including other RPA interacting proteins, and the role of intersegmental transfer. Although the effect of DNA tension on SSB diffusion on DNA has been studied previously (51), diffusive property of yeast RPA on ssDNA longer than 120 oligonucleotides length, or under mechanical tension, has not been reported yet. A recent study by Mersch *et al* have investigated diffusion of human RPA (hRPA) on 20 kbp long ssDNA and reported that the hRPA diffusion does not depend significantly on tension or salt concentration (53).

Diffusion of RPA and SSB has been extensively studied using computation tools (5, 51, 54, 55). We have previously shown force dependent SSB diffusion on ssDNA using coarse grained molecular simulation that aligned with experimental data (51). While previous computational work on RPA diffusion at the small length scale of the ssDNA less than 150 nucleotides provided the molecular details of protein interactions with DNA during diffusion (54–57), they were unable to examine long-range jumps between DNA segments.

In this study, we aimed to understand the diffusion of *Saccharomyces cerevisiae* RPA under various salt conditions and under tension applied across ssDNA. RPA diffuses along ssDNA with one dimensional random walk and its diffusion is reduced with higher ionic strength or with increasing tension. Our findings help us understand how single stranded binding proteins may migrate on single stranded DNA.

## Results

### RPA can diffuse over a long distance on DNA before dissociation

*Saccharomyces cerevisiae* RPA (RPA) was labeled in one of the DBDs with an MB453 fluorophore (58). The fluorophore was positioned either on DBD-A (RPA-DBD-A^MB453^) or DBD-D (RPA-DBD-A^MB453^) using non-canonical amino acids as described (37). To investigate the mechanism of RPA diffusion, we performed confocal fluorescence imaging of fluorescently-labeled RPA on ssDNA under mechanical tension using the LUMICKS C-Trap, an instrument that combines dual optical traps with confocal scanning fluorescence microscopy with single fluorophore sensitivity (59). To create long ssDNA, lambda phage DNA was tethered to two streptavidin beads using biotin on the same strand and was mechanically denatured. Formation of ∼50 kb ssDNA was confirmed by fitting to a Freely Jointed Chain Model (FJC) (60).

Movement of RPA along the ssDNA is recorded as a function of time and visualized as a kymograph (**Figure 1F**). At time zero, each spot corresponds to one RPA molecule bound to the ssDNA ([RPA]=10 pM). Over time, RPA moves back and forth in an apparent diffusive movement, consistent with one dimensional random walk. The displacement (*X*) relative to the initial position is converted into nucleotides using the extension of 48.6 kb ssDNA at a given force (**Figure 1E**). RPA can diffuse over a long distance on DNA before dissociation or photobleaching (**Figure 1F**). Mean square displacement (MSD) plotted vs. time *t* showed an initial linear increase which was fitted using MSD = 2·*D*·*t*(where *D* is diffusion coefficient and *t* is time) to estimate D (**Figure S1**) (61).

### RPA diffusion is affected by DNA tension and salt concentration

We examined the effect of DNA tension on RPA diffusion using two fluorescent versions of RPA labeled at either DBD-A (RPA-DBD-A^MB543^) or DBD-D (RPA-DBD-D^MB543^). We first calculated the diffusion coefficient expressed in μm^2^/s, as measured in the laboratory frame along the direction of force. As shown in Figure 2A and 2B we observed little dependence on force. Next, we accounted for a change in the base per rise (BPR) between nucleotides with force. For example, at 16.2 μM extension at 5 pN, BPR would be 3.4 Å but at higher force with extension of 20 μM, BPR would be 4.1 Å. When we converted the observed displacement into nucleotides then calculated the diffusion coefficient in units of nt^2^/s, we observed a large force dependence (**Figure 2C, 2D**). *D* values were similar for both RPA-DBD-A^MB543^ and RPA-DBD-D^MB543^ and decreased with increasing tension on ssDNA (**Figure 2D, 2E**). For example, for RPA-DBD-D^MB543^, *D* decreased ∼10 fold from 22,000 nt^2^/s at 3 pN to 2,300 nt^2^/s at 28 pN (100 mM KCl) (**Figure 2D**), reminiscent of slower diffusive movements with increasing tension observed for *E. coli* SSB (51). These data also show that conjugation of the MB543 on either DBD-A or DBD-D does not differentially influence their diffusion properties (compare **Figure 2D** and **Figure 2E**), consistent with their similarities observed in the kinetics and thermodynamics of their ssDNA binding properties (27, 37). Thus, we used RPA-DBD-D^MB543^ for further single molecule analysis and will refer to it as RPA^f^.

**Figure 2:**
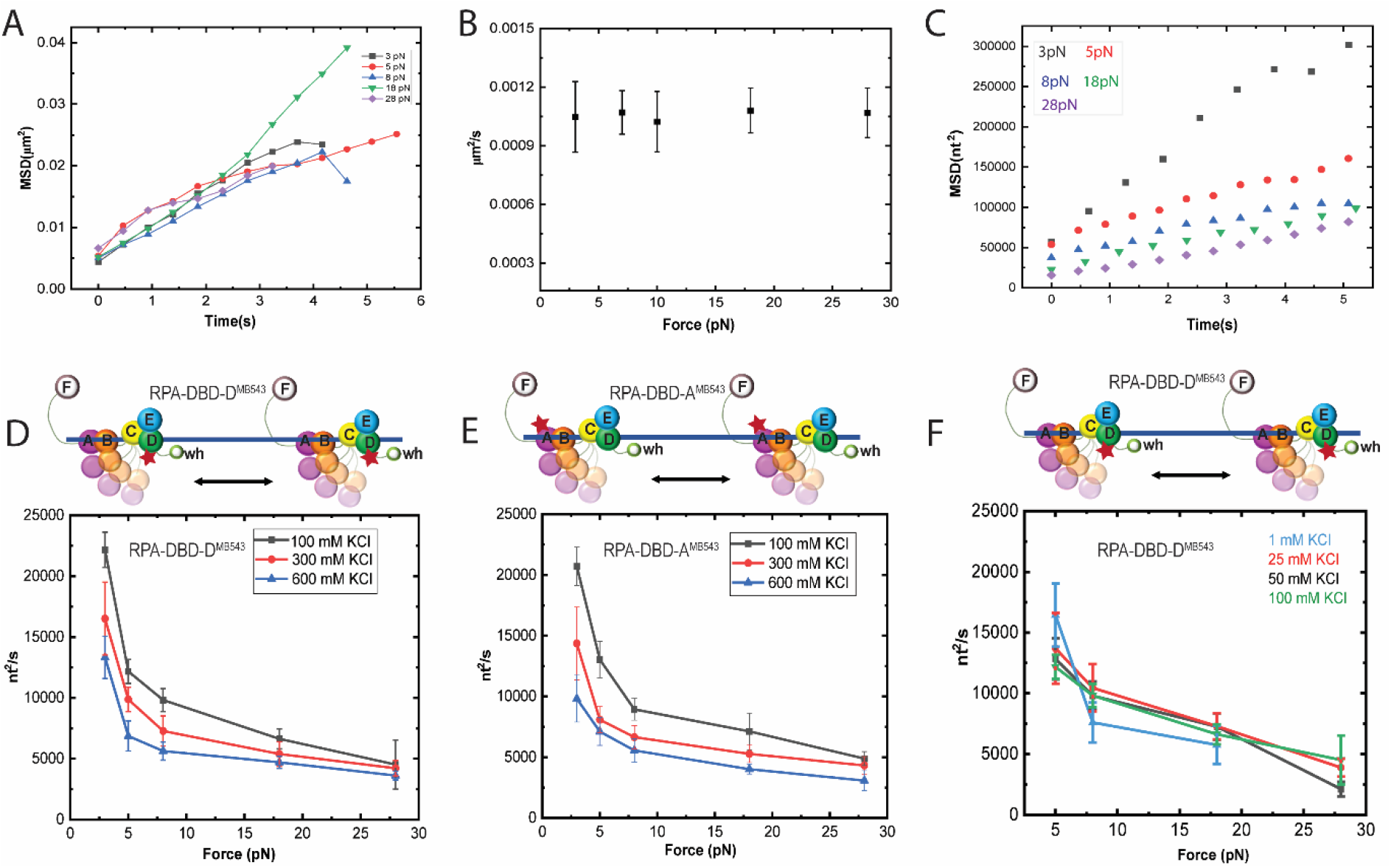
Apparent Diffusion Coefficient of Saccharomyces cerevisiae RPA (RPA) studied as a function of tension on DNA filament and change in salt conditions. A) Mean Squared Displacement (MSD) in micrometer vs time for various forces 3-28 pN. Black 3 pN, Red 5 pN, Blue 8 pN, Green 18 pN, and Purple 28 pN. B) Diffusion coefficient measured as micrometer. C) Mean Squared Displacement (MSD) in nucleotides vs time for various forces 3-28pN. Black 3 pN, red 5pN, blue 8 pN, green 18 pN, and purple 28 pN. D) Diffusion coefficient of RPA protein labelled with MB453 dye at DNA binding domain D (RPA-DBD-D^MB543^) as indicated by red star. E) Diffusion coefficient of full-length RPA protein labelled with MB453 dye at DNA binding domain A (RPA-DBD-A^MB543^) as indicated by red star. F) Diffusion coefficient of RPA-DBD-D^MB543^ at the lower salt concentrations. Experiments were performed at three salt concentrations ranging from 100 - 600 mM KCl while tension was varied from 3 pN to 28 pN. All the error bars are Standard Error of Mean (SEM). Protein concentration is 10 pM with experimental buffer being 100mM KCl (varied), 10mM HEPES pH7.5, 2mM MgCl_2_.

At a low force (3 pN) and high salt (600 mM KCl), RPA^f^ had a diffusion coefficient of 17,000 nt^2^/s (**Figure 2D**) which is approximately 5 times larger than what was previously reported for human RPA at 600 mM KCl (2,800 nt^2^/s) in the absence of tension on 120 nt ssDNA (8). At this diffusion coefficient, an RPA can migrate over about 200 nt in 1 second. At forces higher than 18 pN, the diffusion coefficient for yeast RPA^f^ decreases to ∼3000 nt^2^/s at 600 mM KCl which is comparable to that of human RPA diffusion on short DNA at zero force (**Figure 2D, 2E**). One possible explanation is that for long ssDNA, RPA may use intersegmental transfer between distant sites, a process that would be hindered when the DNA is stretched mechanically.

Next, we looked at the effect of ionic strength. RPA became less mobile with increasing salt concentration for all force values tested (**Figure 2D, 2E**). For example, *D* for RPA^f^ decreased by a factor of three when [KCl] increased from 100 mM to 600 mM (**Figure 2D, 2E**). Salt concentration influences the way various combinations of DBDs associate with ssDNA, with higher salt promoting binding of more DBDs (8, 32, 36). If more DBDs bind to the DNA, RPA may diffuse slower due to an enhanced number of contacts.

RPA switches from the 22 nt binding mode to the 30 nt binding mode when salt concentration increases from 100 mM to 600 mM, but the binding mode stays at 22 nt below 100 mM down to 1 mM (8, 36). Therefore, we performed the experiment under low salt concentrations to further test if the effect of salt concentration on RPA diffusion is indeed due to changes in binding mode. We did not see a significant change in *D* between 1 mM and 100 mM KCl (**Figure 2F**). Therefore, our data are consistent with the model where RPA is more mobile in the 22 nt binding mode compared to the 30 nt binding mode.

### Coarse grained simulations of RPA diffusion

To understand the effect of force and salt on the diffusion of RPA on ssDNA in molecular details, we used coarse-grained molecular simulations. We started by using different starting conformations of a fixed length ssDNA, where the end-to-end distances were different. Each of the systems were simulated keeping the ends of the DNA fixed in space, while the other parts of the DNA and the protein were allowed to move freely. This represents the experimental scheme for optical trap where traps were fixed at a specific distance with ssDNA held between two traps. Tuning of the end-to-end distance effectively control the force on the DNA where higher end-to-end distance represents higher force on ssDNA (**Figure S2**).

A representative simulation of RPA on 860 nt ssDNA (0.5 pN force, 30 mM salt) is shown as five snapshots of RPA-bound ssDNA’s conformation. In snapshot 2, RPA bridges two distant sites that are separated by 168 nt (**Figure 3A**). Around this time point, the index of the nucleotide closest to the protein center of mass (COM) changes rapidly back and forth by Δnt = 168, eventually resulting in a large jump in position by that amount (**Figure S3**). We proposed that the looped state, with the loop size of 168 nt, is an intermediate for intersegmental transfer. Snapshot 4 shows another looped state of a 51 nt loop, resulting in a smaller scale intersegmental transfer. The size of intersegmental transfer is widely distributed and became progressively smaller as the force increased (**Figure 3B**). At high force, the DNA is more stretched, which hinders the formation of a large ssDNA loop thereby reducing the probability of large intersegmental transfer events. Indeed, the wait time between any two consecutive intersegmental transfers with Δnt > 10 dramatically increases as the force increase (**Figure 3C**). A similar force dependence was observed also for high salt simulations (1000 mM) (**Figure S4D, S4E** and **S4F**) with the main difference being that in higher salt, the intersegmental transfer is often shorter in jump size and their frequency is lower, consistent with lower mobility in high salt we observed experimentally (**Figure 2**). The diffusion coefficient *D* calculated from the simulated trajectories also showed salt and force dependence (**Figure 3D**). In high salt, a 15-fold increase of the force resulted in a ∼ 6 folds decrease in the *D* value, analogous to a 9-fold increase in force causing a ∼4-fold decrease in *D* observed experimentally (**Figure2, 3D**). At the low salt simulation, *D* was 3-4 times greater than in the high salt simulation, similarly to the fractional change in *D* determined experimentally (**Figures 2, 3D**).

**Figure 3:**
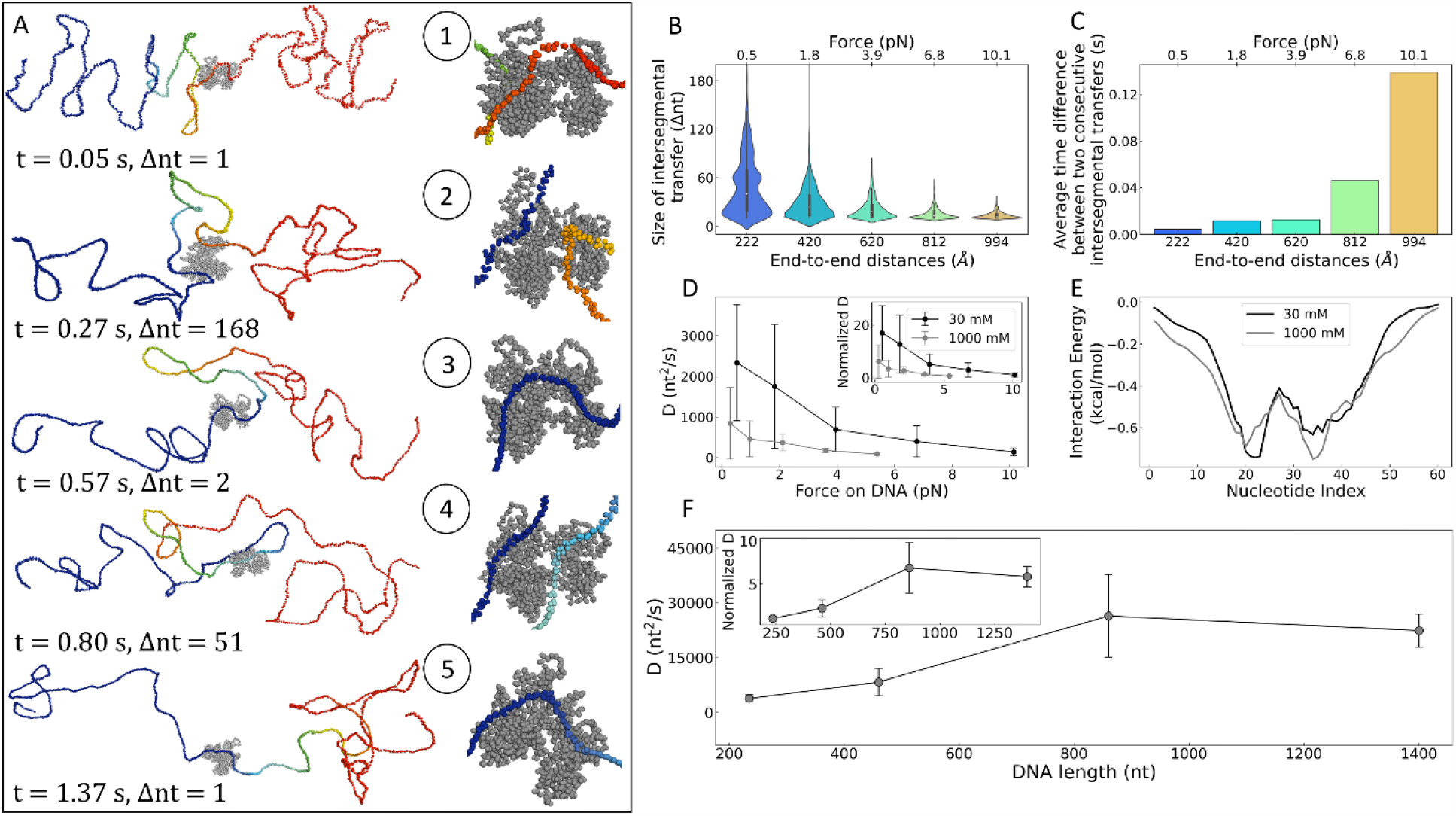
Diffusion of RPA on ssDNA through intersegmental transfer. A) Five representative snapshots of RPA-ssDNA binding sampled variation of the nucleotide index (nt) closest to the centre-of-mass (COM) of the RPA protein is shown throughout a representative simulation trajectory for ssDNA of length of 860 nucleotides ssDNA with a force of 0.5 pN on the ssDNA (which corresponds to an end-to-end distance of 817 Å. B-C) Characteristics of the intersegmental transfer events of RPA when diffusion along ssDNA under force applied on its ends. These simulations were performed at 30 mM. B) Violin plots of the length of ssDNA, Δnt, skipped in intersegmental transfer of RPA when diffusing along ssDNA at five different forces applied on its two ends [the applied force (shown in the upper X axis) results with end-to-end distance varying between 222 - 994 Å, (shown in the lower X axis)]. The simulations at different force applied on the two ssDNA ends are shown with different color. C) The kinetics of the intersegmental transfer is indicated by the average time difference between any two consecutive intersegmental transfer events (having length Δnt > 10) for the five systems at 30mM.D-E) Variation of diffusion coefficient at different force and salt concentrations. D) Variations of diffusion coefficient for diffusion of RPA along ssDNA for different strength of force applied on its two ends, at low (black) and high (grey) salt concentrations. The inset shows the diffusion coefficients normalized to the value at high applied force. The error bars represent the standard deviation in the diffusion coefficient obtained from 30 identical simulations. E) The potential energy of the interaction between RPA and ssDNA of 60 nucleotides as a function of the ssDNA nucleotide index for low and high salt concentrations. A ssDNA of 60 nucleotides have been utilized to focus on the protein-DNA interaction at the binding site as per the RPA-ssDNA crystal structure. F) Variation of RPA diffusion coefficient as a function of ssDNA length. The inset shows the normalized diffusion coefficient relative to the diffusion coefficients for the shortest simulated ssDNA. The error bars represent the standard deviation of the diffusion coefficients obtained from 30 independent trajectories.

To gain insights into the molecular origin of the salt effect, we calculated the binding energy vs nucleotide position at the binding interface RPA-ssDNA for 235 nt long ssDNA (**Figure 3E)**. For most of the regions of the protein-ssDNA interface, the interaction is more stable (i.e., more negative interaction energy) at higher salt concentration. The tighter interaction between RPA and ssDNA at high salt is consistent with the higher binding mode. We do not completely understand the nature of contacts between the domains of RPA as salt could also influence such interactions and add another layer of complexity in deciphering these transitions.

The ssDNA used in the experimental study is much longer than the 235 nt ssDNA used in the simulations thus far. To explore the effect of the ssDNA length on the characteristics of intersegmental transfer, we performed simulations with different ssDNA lengths (235, 460, 860 and 1400 nucleotides). As the DNA length increases from 235 nucleotides to 860 nucleotides, there is an increase in the diffusion coefficient *D* (**Figure 3F**), likely due to higher probability of large size intersegmental transfer events (**Figure 3B, S4D)**. Beyond 860 nucleotides, the change in the *D* value was negligible (**Figure 3F**). The saturation of the intersegmental transfer events is also reflected in the size distribution and occurrence probability of intersegmental transfer (**Figure S5B, S5C**). This result suggests that for a better comparison to experimental results we should use simulations obtained using the longer ssDNA (860 nt and 1400 nt). They showed a 10-fold increase in the *D* values (**Figure 3G**, inset plot) compared to 235 nt long DNA, suggesting that *D* values with the short ssDNA should be multiplied by factor of by 10 to compare to the experiments. Indeed, we obtained a diffusion coefficient ∼ 2500 nt^2^/s with 235 nucleotides DNA (**Figure 3F**), while in experiments for a 48.5 kbp ssDNA the measured *D* value obtained was ∼ 10 times higher (**Figure 2D, 2E**).

### The ‘dynamic half’ of RPA consisting of OB-F, DBD-A, and DBD-B has higher mobility

Diffusion of RPA is proposed to occur through dynamic binding and rearrangements of the individual DBDs (27, 37).These domains were recently classified to function as two halves: “FAB”, composed of OB-F, DBD-A, and DBD-B, is considered ‘dynamic’ based on their higher propensity to detach from ssDNA. The “trimerization core” (Tri-C), composed of DBD-C, DBD-D and DBD-E, is considered ‘less-dynamic’, contributing more to the stability of RPA-ssDNA interactions (27, 33, 37). FAB, lacking the Tri-C, does not show significant DNA dependent protection from proteolysis, likely because its interaction with DNA is not as stable as the fulllength protein, frequently exposing the protein surface (27). The occlusion size of FAB on ssDNA is proposed to be ∼8 nt (8, 28). To test the contributions of these two halves to RPA diffusion, we measured the diffusion properties of FAB. FAB is highly diffusive with diffusion coefficient of 210,000 nt^2^/s at 5 pN which is about 15 times larger compared to the full-length RPA constructs under same tension (**Figure 4**). FAB binding to DNA is more transient, with its average residence time on DNA of 18 seconds before dissociation compared to the less diffusive WT RPA with an average residence time of one minute (**Figure 4C, 4D, 4E**). The larger diffusion coefficient for FAB is likely due to fewer nucleotide contacts with ssDNA, also causing fast dissociation. FAB also showed similar salt and tension dependence of diffusion coefficient as in the full-length proteins (**Figure 4A, Figure 2**). Unfortunately, such experiments with TriC (DBD-C, RPA32, and RPA 14) alone were not possible due to the instability of the complex when isolated.

**Figure 4:**
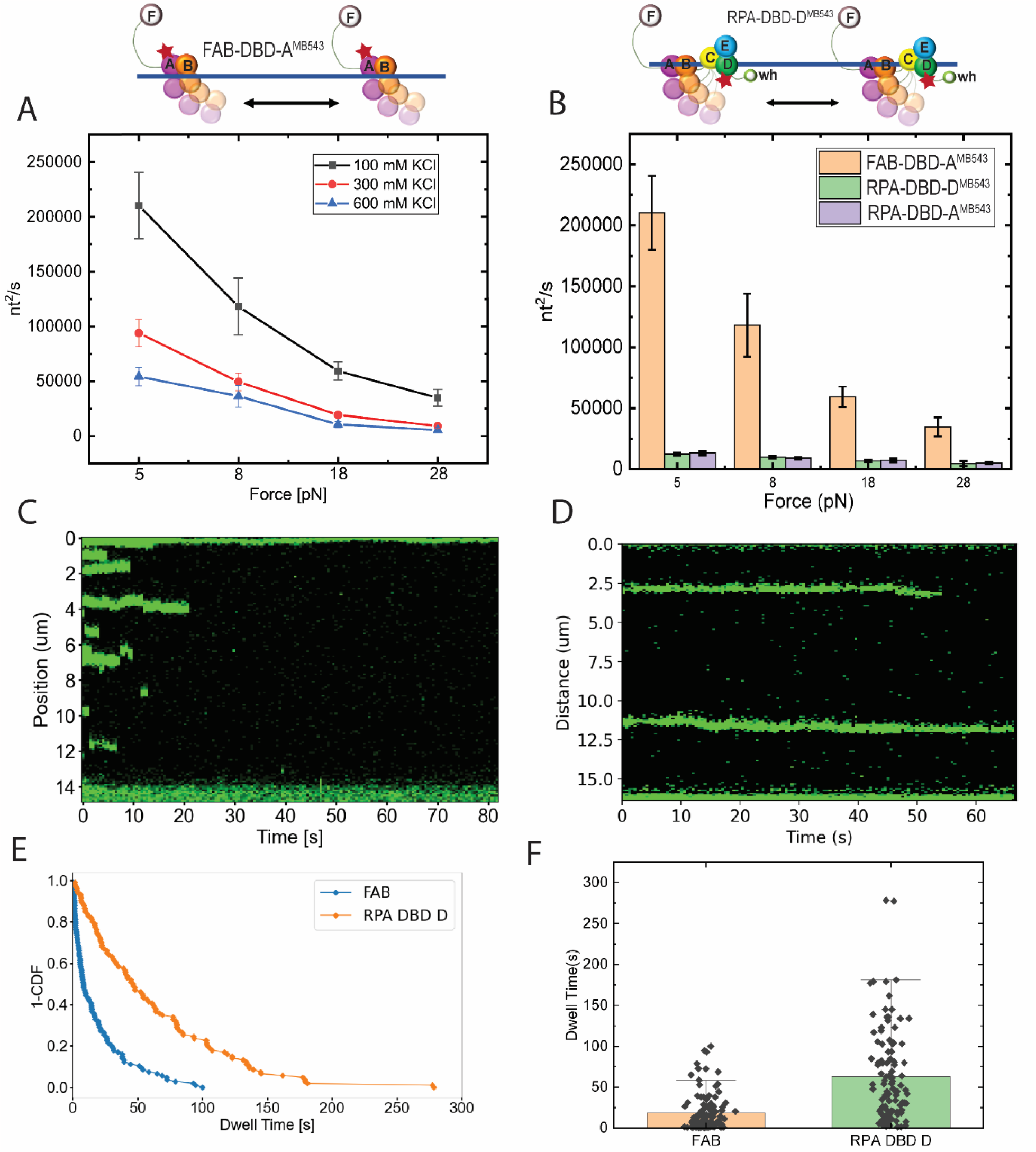
Apparent Diffusion Coefficient of saccharomyces cerevisiae RPA (RPA) studied as a function of tension on DNA filament and change in salt conditions. A) Diffusion of truncated protein with only FAB domains from RPA70. B) Comparison of diffusion coefficient of four different variants of RPA studied. RPA DBD A and RPA DBD D data are taken as in figure 2 while FAB data is taken from Figure 4 A. The Schematic of RPA shown on top is a representative RPA diffusion where RPA is labeled at DNA binding domain D (RPA-DBD-D^MB543^) as in Figure 2D.C) A typical kymograph for FAB. FAB is very diffusive in short range and dissociates from DNA. D)A typical kymograph of full-length RPA. E) A 1-CDF plot depicting how fast FAB dissociates from DNA compared to full length RPA.F) Dwell time of RPA full length and FAB on ssDNA. Salt conditions ranged from 100 mM KCL to 300 mM KCL while tension ranged from 3pN to 28pN with error bars being SEM. FAB concentration was 1nM.Other buffer components were 10mM HEPES pH7.5,2mM MgCl_2_.

### RPA diffusion under RPA crowding

Next, we looked at the effect of excess unlabeled RPA WT (full length RPA with no label in any DBD) in the diffusional properties of RPA^f^ (**Figure 5A**) in order to mimic the crowding effect provided by additional RPA proteins bound to the same DNA (37, 62). At unlabeled RPA WT to RPA^f^ ratio of 15 or higher, *D* decreased measurably. *D* dropped from 13,000 nt^2^/s in the absence of excess RPA to 900 nt^2^/s when 100 times excess unlabeled RPA was available, probably because the available free space is constrained (**Figure 5**). As the concentration of RPA was increased, the effective distance traversed by a single RPA decreased; for example, RPA diffused up to 200 nts for WT to RPA^f^ ratio of 30 compared to up to ∼2000 nts for WT to RPA^f^ ratio of 1 likely due to collisions with other RPA molecules consistent with the decrease in diffusion coefficient (**Figure 5B, 5C**).

**Figure 5.**
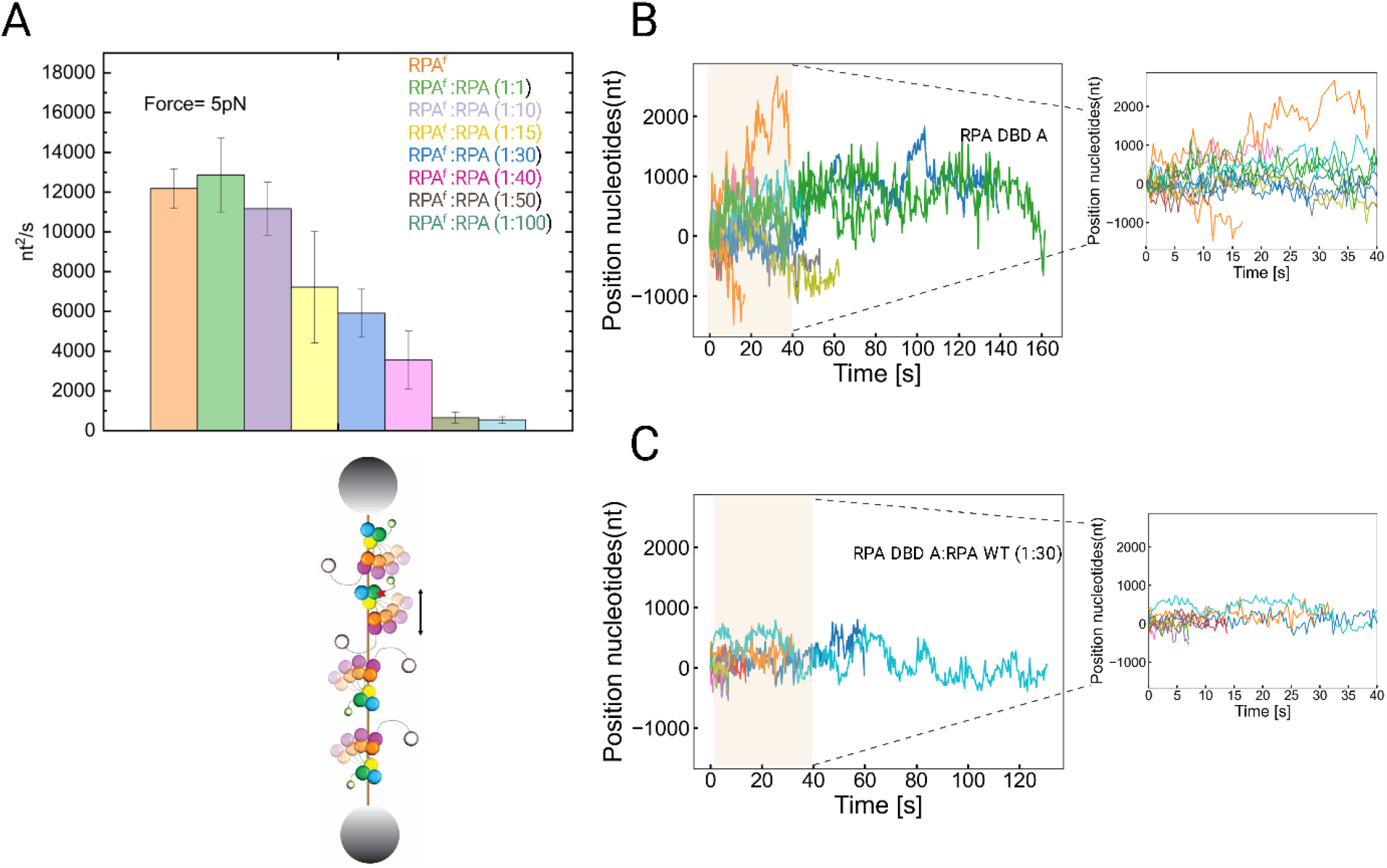
RPA diffusion under the influence of free unlabeled RPA and chaperone Rtt105. A) Apparent diffusion coefficient of RPA DBD D (RPA labeled at DBD D as indicated by red star) under the presence of various concentrations of unlabeled RPA WT. Desired ratio of proteins were applied to the same microfluidic channel by incubating together for 1-2 min on ice. Error bars are SEM. B) Distance traversed by RPA under the condition where there are no collisions with other RPA molecules. C) Distance traversed by RPA^f^ where WT to RPA^f^ ratio of 30 was used.

## Discussion

Using optical trap combined with single molecule confocal fluorescence microscopy, we directly observed the diffusional movements of RPA along mechanically stretched ssDNA in real time. The diffusion coefficient is 21000 nt^2^/s at 3 pN at physiological salt concentration of 100 mM KCl. Coarse grained simulations provide evidence for the importance of intersegmental transfer.

We observed a remarkable decrease in diffusion coefficient with increasing salt concentration. RPA can adopt different ssDNA binding modes on ssDNA as per the ionic condition. Nguyen *et al* (8) showed that as salt concentration increases, the occlusion size, the number of nucleotide contacts made by human RPA on ssDNA, increases. Kumaran *et al* proposed a similar binding mode switch for yeast RPA, a lower DNA binding mode of 18 nt in low salt and a higher DNA binding mode of 28 nt in high salt (36). The decrease in *D* we observed when KCl concentration increased from 100 mM to 600 mM could be a result of such switch in binding mode (8, 36). The resulting depletion of DNA binding sites on the RPA surface would hinder the formation of a transient bridge to a distant site on DNA, reducing RPA mobility. Consistent with this explanation, we see no difference in *D* in salt concentration ranging from 1 mM to 100 mM (**Figure 2F**), the range for which Nguyen *et al* did not see any change in binding mode.

Persistence length of ssDNA decreases with increasing salt (63), making ssDNA more flexible through screening electrostatic repulsion between negatively charged backbone phosphates (64, 65). Higher flexibility may make it possible to obtain tighter contact with DNA binding sites spread over the RPA surface. Indeed, our molecular dynamics simulations based on the crystal structure of RPA-ssDNA complex (29) suggest that the higher salt concentration induces stronger interaction (**Figure 3E**). Overall, the salt effect on diffusion of RPA could be governed by the binding mode of RPA, ssDNA flexibility and RPA-ssDNA interaction strength. Additionally, recent cross-linking and hydrogen-deuterium mass spectrometry experiments reveal extensive interactions between the DBDs and PIDs of RPA(27, 66, 67). Thus, additional salt-induced changes to such intra-RPA contacts could also influence the diffusion properties.

An earlier report of force dependent diffusion of ssDNA binding protein SSB (51) showed that the diffusion coefficient (*D*) dropped by a factor of 6 as the force was increased from 3 pN to 5 pN (51). Because SSB dissociated from ssDNA at ∼8 pN, the range of tension examined was small (51, 68–70). The DNA construct used for the SSB study could not form secondary structure as ssDNA were generated using rolling circle amplifications of a short sequence devoid of secondary structure. In the current study of yeast RPA, while diffusion was slower than SSB, it was higher than the values reported for human RPA by Nguyen *et al* (8): i.e. 17000 nt^2^ /s in our study vs 2800 nt^2^/s in theirs, both in 600 mM KCl. They also used 120 nt oligos with no secondary structure possibilities (8). Our experiments performed using lambda DNA would allow some secondary structures to form at lower forces, potentially making intersegmental transfer more effective.

The size of intersegmental transfer also decreases with increasing force as we show through molecular simulations (**Figure 3B**). In the absence of force, DBDs of RPA can bind distant sites at the same time and effectively stabilize the condensed DNA (65). Recent study on human RPA did not observe a significant influence of force on its diffusion (53). Our study on yeast RPA showed a clear force dependence on the diffusion of the protein both in experiment and simulations. Although human RPA and yeast RPA are structurally similar, experimental observations have suggested that yeast RPA is more dynamic than human RPA (37, 40, 71, 72). This could also explain the differential effect of force on diffusion of human RPA and yeast RPA (55, 73). We note that the diffusion coefficient expressed in μm^2^/s, that is in the distance measured in the laboratory frame along the direction of force, showed little dependence on force (**Figure 2B**). It is when we convert that distance to the number of nucleotides that we see a strong force dependence (**Figure 2D**).

Another possible explanation for force dependent decrease in diffusion can be the reptation mechanism. Reptation involves DNA bulge formation on the RPA surface, storing 1–7 nt of ssDNA involving aromatic interactions between protein and ssDNA. Bulge dissolution *i*.*e*. release of 1-7 nt in the bulge leads to RPA diffusion on ssDNA (5, 54, 55). There has not been direct experimental evidence for this mechanism in the case of RPA, but bulge formation has been reported as a possible diffusion mechanism for SSB in fluorescence-force spectroscopy analysis (74) and also in molecular simulation studies of RPA (55). As we increase the tension on ssDNA, the possibility of RPA forming a bulge will be diminished as ssDNA is stretched. Conversely, under low tension where relaxed DNA is present, RPA can diffuse faster with bulge formation increasing the diffusion.

We found that when high density of RPA is present on ssDNA, 1D random motion of individual RPA molecules becomes restricted, leading to an overall decrease in diffusion coefficient (**Figure 5**). Whether the crowing effect mainly acts through inhibiting local migration in small, a few nt steps, as is expected for reptation, or by hindering larger scale movements such as intersegmental transfer is presently unknown.

Our study suggests that full length RPA is less dynamic than FAB, a truncated RPA that lacks the trimeric core. FAB was 15 times more mobile than full length RPA (**Figure 2D, 4A**). Further studies are needed to determine if the higher mobility is due to an increase in size or frequency of intersegmental transfer.

## Methods

### Protein Purification

Replication Protein A was purified and labeled as detailed in (38, 58, 73).

### Optical Trap combined with confocal microscopy

48.5 kbp Lamda DNA construct was prepared with three biotins on either end of DNA(60). Lambda DNA was purchased from Roche Inc. Short oligos to anneal to the sticky ends were purchased from Integrated DNA Technologies Inc. DNA were stored in TE buffer (10 mM Tris-HCl, pH 8.0 and 0.1 mM EDTA). Optical trap experiments were performed using a commercial dual optical trap combined with confocal microscopy and microfluidics [C-trap] from Lumicks BV Inc. The microfluidic chamber was passivated at the start of every experiment day using the protocol: 0.1% BSA (Sigma) was flowed at 1 bar for 5 minutes then at 0.4 bar pressure for 25 minutes, followed by a 10-minute rinse with Milli-Q water at 1 bar pressure for 5 min and 0.4 bar pressure for 5 minutes, followed by 0.5% Pluronic F-127 flowed at 1 bar pressure for 5 min then 0.4 bar pressure for 25-minutes, followed 1 bar pressure for 5 min and 0.4 bar pressure for 5 minutes with 1X PBS.

Streptavidin coated polystyrene particle beads of average size 4.8 μM [0.5 % w/v] (Spherotech Inc.) were diluted 1:250 in 1X Phosphate Saline Buffer (PBS) that contains 137 mM NaCl, 2.7 mM KCl, 8 mM Na_2_ HPO_4_, and 2 mM KH_2_PO_4_(10X buffer purchased from Thermofisher Scientific USA) and 1-2 nM of DNA were made in 1X PBS. DNA was captured between two streptavidin beads and mechanically denatured by moving one bead to the overstretched region to create ssDNA. ssDNA was confirmed by fitting force-distance (FD) curve to Freely Jointed Chain model [FJC] (contour length 48.5 kbp / 27.160 um; persistence length 0.9 nm; stretch modulus 1000 pN) in real time(75, 76). DNA was held for 5 s in the fully ssDNA state then returned to a required tension on ssDNA position for the fluorescence experiments.

RPA-DBD-D-MB543, RPA-DBD-A-MB543, and RPA WT were stored at storage buffer (30 mM HEPES pH 7.8, 200 mM KCl, 0.02 % Tween-20, 10 % glycerol, and 0.2 mM EDTA pH 8.0) and diluted to 1 nM with experimental buffer (30 mM HEPES pH 7.8, 100 mM KCl, 6 % Glycerol, 5 mM MgCl_2_) just before the experiment then 10 pM final concentration of RPA was used during the experiments. Imaging buffer 0.8 % (w/v) dextrose, 165 U/mL glucose oxidase, 2170 U/mL catalase, and 2-3 mM Trolox was used to increase the fluorescence lifetime of the fluorophores. Imaging settings were 2-3 ms exposure time (per pixel), red excitation 638 nm, and green excitation 561 nm. Pixel size was kept constant at 100nm. Excitation wavelengths were Green (575-625 nm), Red (670-730nm). Laser power was selected to maintain the 5 micro-watt power at the objective. Note that our experiments required pixel time of 3 ms to capture a single RPA molecule. Reducing the pixel time would often require higher RPA concentration of protein where less bright RPA could not be captured but will still contribute to the diffusion of RPA being observed.

### DNA Oligos

Lambda DNA (Purchased from Roche)

Oligo 1: GGG CGG CGA CCT GGA CAA

oligo 2: AGG TCG CCG CCC TTT TTT T/iBiodT/T /iBiodT/T/iBiodT/

oligo 3:/iBiodT/T/iBiodT/ T/iBiodT/T TTT TTT AGA GTA CTG TAC GAT CTA GCA TCA ATC TTG TCC

### Data Analysis and Sharing

Data was analyzed using custom python script detailed in Pylake API from Lumicks (77). Python version 3.8.6 is used. Origin pro 2021b was also used. Some illustrations were made using Biorender.com. Similarly, all the codes are available in Github(https://github.com/spangeni/RPA-Diffusion.git). All the data has been included in the manuscript. Raw data will be made available upon reasonable request.

### Simulation Methods

#### 1. Generating protein coarse-grained model

To investigate the interaction of RPA with ssDNA at the molecular level, a coarse-grained model was used. In this model, each of the protein residues are represented by two beads placed at the C_α_ and C_β_ positions. For the charged amino acids (K, R, H, D and E), respective charges were placed at the C_β_ position. Simulations have been performed using a native topology-based model which included nonspecific electrostatics (non-native) interactions and used Lennard-Jones potential to represent native contact interactions. For protein modelling, an approach similar to that described in Refs. (54, 56) have been utilized, where the internal energy of the protein is designated by E_prot_, where,

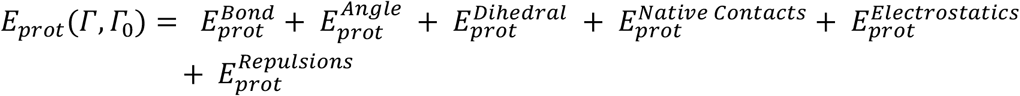

where Γ denotes a particular conformation during the simulation trajectory and Г_0_ denotes the native conformation. The potential energy of any conformation during the simulation trajectory consists of the following terms,

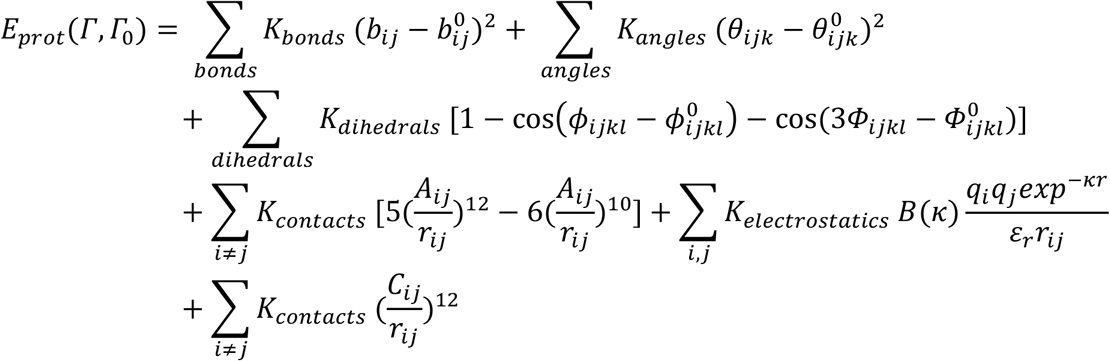

where K_bonds_, K_angles_, K_dihedrals_, K_contacts_, and K_repulsion_ = 100 Kcal mol^−1^ Å^−2^, 20 Kcal mol^−1^, 1 Kcal mol^−1^, 1 Kcal mol^−1^, and 1 Kcal mol^−1^, respectively. Additionally, b_ij_ is the distance between bonded beads i and j in Å, and b^0^_ij_ is the optimal distance between bonded beads i and j in angstroms; θ_ijk_ is the angle between subsequently bonded beads i–k in radians, θ^0^_ijk_ is the optimal angle between subsequently bonded beads i–k in radians, *ϕ*_ijkl_ is the dihedral angle between subsequently bonded backbone beads i–l in radians, and ϕ^0^_ijkl_ is the optimal dihedral angle between subsequently bonded backbone beads i–l in radians. A_ij_ is the optimal distance between beads i and j in contact in angstroms, and r_ij_ is the distance between beads i and j in angstroms in a given conformation along the trajectory. Optimal values were calculated from the atomic coordinates of the relevant Protein Data Bank (PDB) structure file. C_ij_ is the sum of radii for any two beads not forming a native contact; the repulsion radius of the backbone bead (C_α_) was 1.9 Å, and the radius of the side-chain bead (C_β_) was set to 1.5 Å. The electrostatic interactions were modeled by the Debye–Hückel potential, and parameters from previous studies (54, 56) have been used. The simulations were run at a temperature where protein remains folded and fluctuate around the native state.

#### 2. Generating ssDNA coarse-grained model

In the coarse-grained model of the single-stranded DNA (ssDNA)(78) each of the nucleotide was represented by three beads, positioned at the geometric centre of phosphate (P), sugar (S) and base (B); while the S and B bead were neutrally charged, the P bead beard a negative charge. The internal potential energy of the ssDNA was designated by E_ssDNA_, which is given by the following expression,

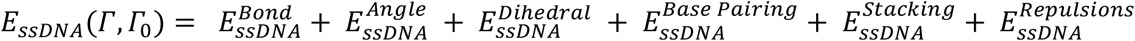

In this expression, the term 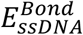 represents the contribution from the backbone consisting of covalently linked phosphate (P) and sugar (S), also, covalently linked sugar (S) and 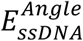 ssDNA is the potential for bond angles that is applied between all the following three neighboring beads: (P_i_-S_i_-B_i_), (B_i_-S_i_-P_i + 1_), (P_i_-S_i_-P_i + 1_), and (S_i_-P_i + 1_-S_i + 1_) with K_angles_ = 20. 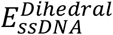 consists of two kinds of dihedral potentials. One is the potential of the dihedral angle formed by the beads B_i_, S_i_, S_i + 1_, and B_i + 1_ with K_Dihedral_ = 0.5. The other is the potential from the dihedral angle formed by four consecutive phosphate beads (P_i_, P_i+1_, P_i+2_ and P_i+3_); and this K_Dihedral_ is either 0.7 or 0 which makes the persistence length of the DNA 32 and 17 Å respectively.

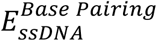 arises from the contribution of the interaction between complementary base-pairs, which is absent in case of ssDNA, so 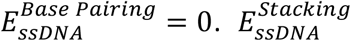 arises from the interaction between two consecutive bases, B_i_ and B _i + 1_ and can be expressed as 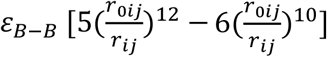, where *r*_0*ij*_is the typical distance between two consecutive bases (i.e., 3.6 Å) and strength of interaction ε_*B*−*B*_ depends upon the identity of bases (i.e., whether A, T, C or G). The expression for the repulsion term, 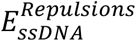 is similar to the repulsion term in case of proteins, only the radii of the base, phosphate, and sugar are 1.5, 3.7, and 3.7 Å respectively and the repulsion is applied between two beads which satisfies the condition |i – j| > 6.

#### 3. Modelling protein-ssDNA interactions

The interaction energy between protein and single stranded DNA is given by,

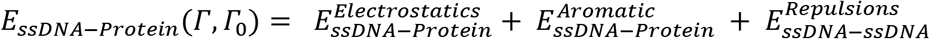

The repulsion is applied between all beads of the protein and all beads of the ssDNA. The electrostatic interactions acting between all the charged beads in the system are modeled by the Debye–Hückel potential. These interactions are nonspecific, and the phosphate groups of the ssDNA can interact with any charged residue in the protein. The aromatic interaction, like base stacking, is modeled by the L-J potential, with a base–aromatic amino acid interaction strength of ε_*B*−*AA*_. The parameter ε_*B*−*AA*_= 0 for all the nonaromatic amino acids, and ε_*B*−*AA*_> 0, only for aromatic amino acids (W, F, Y, and H). The strength of interaction ε_*B*−*AA*_ for aromatic amino acids depends on their identities.

#### 4. Modelling force

Next, to understand the effect of force on ssDNA in molecular details, we started with the RPA-ssDNA coarse-grained model obtained from the original crystal structure (PDB ID; 4GNX). Since, in the crystal structure only 29 nucleotide long ssDNA was present, we modelled the ssDNA as longer one (initially 235 nucleotides) by keeping the RPA-bound fragment of 29 nucleotides in the middle and identical to the original crystal structure. Now, after generating the coarsegrained model, pulling force was applied to both ends of the DNA to obtain different initial structures having different end-to-end distances representing different forces. To that end, five different end-to-end distances were utilized, namely, 222, 420, 620, 812 and 994 Å. This end-to-end distances were converted to respective force values using the FJC model (79), which resulted in forces 0.5, 1.8, 3.9, 6.8 and 10.1 pN respectively on the ssDNA.

Now, with these five initial structures, molecular dynamics simulation was performed keeping both ends of the ssDNA fixed in space while the protein and the rest of the DNA was allowed to move freely. The temperature was 0.4 (arbitrary units) and the salt concentration was kept at 0.065 mM during the simulation, and the simulations were run 5×10^7^ simulation steps, saving the coordinates at each 1000 steps. The simulation temperature is chosen in such a way that the protein remains folded during the simulation and fluctuates around its native state. Importantly, the salt concentration mentioned before, regulates the electrostatics interaction between the protein and DNA, but since in our model, no electrostatic interaction is present between the DNA beads, this salt concentration cannot confer flexibility or rigidity to the DNA molecule which is sensitive to the salt concentration.

#### 5. Modelling effect of salt

To model the effect of salt in the RPA-ssDNA interactions, we modified the persistence length of the DNA that is salt dependent. From previous studies (56, 64), it is well known that, as the salt concentration is increased, the ssDNA becomes more flexible and its persistence length decreases. Also, in a similar coarse-grained model of ssDNA, the persistence length was modified by changing the dihedral constant K_Φ_ (56). In our default model, the dihedral constant was 0.7, which resulted in a persistence length of 32 Å (56). As per the experimental studies by Murphy et al, this persistence length of the ssDNA is equivalent to that in the 30 mM salt (56, 64). Thus, to increase the DNA flexibility, which would correspond to higher salt concentration, we modified the strength of the dihedral constant K_Φ_ to 0, which resulted in a persistence length of 17Å, the flexibility of which would correspond to that in the salt concentration 1000 mM. It is noteworthy that, since the force-extension relationship according to the FJC model explicitly depends upon the persistence length of the DNA, the conversion of the end-to-end distances to force between the two ends depends on the salt concentration. Therefore, the end-to-end distances of 222, 420, 620, 812 and 994 Å correspond to forces of 0.5, 1.8, 3.9, 6.8 and 10.1 pN for salt concentration of 30 mM (K_Φ_ = 0.7) and to values of 0.3, 1.0, 2.1, 3.6 and 5.4 pN for salt concentration of 1000 mM (i.e., K_Φ_ = 0).

#### 6. Effect of DNA length

As mentioned previously, the ssDNA in our simulations is of length of 235 nucleotides. Since the ssDNA in the experiments is much longer, which may affect the probability of intersegmental transfer, we simulated the diffusion of RPA in several systems for varying length of ssDNA. We used ssDNA of length of 235, 460, 860 and 1400 nucleotides. For these four systems, the end-to-end distances were kept as 222, 435, 817 and 1320 Å, respectively, which corresponds to similar force values of ∼ 0.5 pN, for each case.

#### 7. Calculation of diffusion coefficient and intersegmental transfer

The diffusion of RPA along ssDNA is characterized by the diffusion coefficient. In order to do that, first, the distance of all the ssDNA nucleotides were calculated from the protein centre of mass (COM) for every saved trajectory throughout the simulation. The nucleotide index closest to the protein COM in each step was identified as M. Now, the three-dimensional diffusion coefficient (D) was calculated from the mean square displacement (MSD) of the centre-of-mass of RPA moving along the ssDNA:

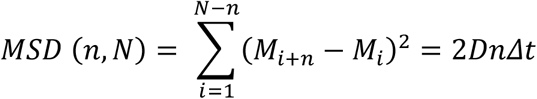

where *N* is the trajectory length in time steps, n is the measurement window ranging from 1 to *N, Δt* is the time step interval, and *M* is the nucleotide index closest to the protein COM at the corresponding time step. The diffusion coefficient was calculated between time frames 200 to 800 for diffusion on DNA.

Further, to convert the simulation time steps in real time unit, we took a small fragment of the ssDNA consisting of 20 nucleotides to monitor its dynamics. The coarse-grained model of this 20-nucleotide ssDNA was simulated for 1 × 10^6^ simulation time steps allowing the whole ssDNA to move freely in space. We ran 50 copies of the same simulation to obtain the lifetime for forming a closed circular ssDNA (i.e., formation of a contacts between the two ends) in the units of simulation time steps. This lifetime was matched with the experimental lifetime in real units in seconds obtained from the experiments by Kim et al (80), and we obtained the equivalent time units in seconds for a simulation time step. We found that each simulation time step corresponds to 3.267 × 10^-8^ s. Thus, the diffusion coefficient obtained in the units of nt^2^/timesteps were then converted to nt^2^/s.

To calculate the size of the intersegmental transfers (Δ*nt*, which corresponds to the length of the ssDNA bridged during the jump), the absolute value of the difference of nucleotide index (M) closest to protein COM between two consecutive saved trajectories (specifically 1000 consecutive simulation steps) were calculated which can be expressed as follows,

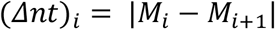

where, M_*i*_ represents the nucleotide index closest to the protein COM at *i*^th^ simulation time step, while M_*i*+1_ stands for the nucleotide index closest to the protein COM at *i*+1^th^ simulation time step. We define this difference of nucleotide index as intersegmental transfer if *Δnt* > 10.

## Acknowledgements

We would like to thank the team at Lumicks for their technical help. Momcilo Gavrilov for initial C-trap training to the author. This article is subject to HHMI’s Open Access to Publications policy. HHMI lab heads have previously granted a nonexclusive CC BY 4.0 license to the public and a sublicensable license to HHMI in their research articles. Pursuant to those licenses, the author-accepted manuscript of this article can be made freely available under a CC BY 4.0 license immediately upon publication.

