## supplementary file for "Rapid long-distance migration of RPA on single stranded DNA occurs through intersegmental transfer utilizing multivalent interactions"

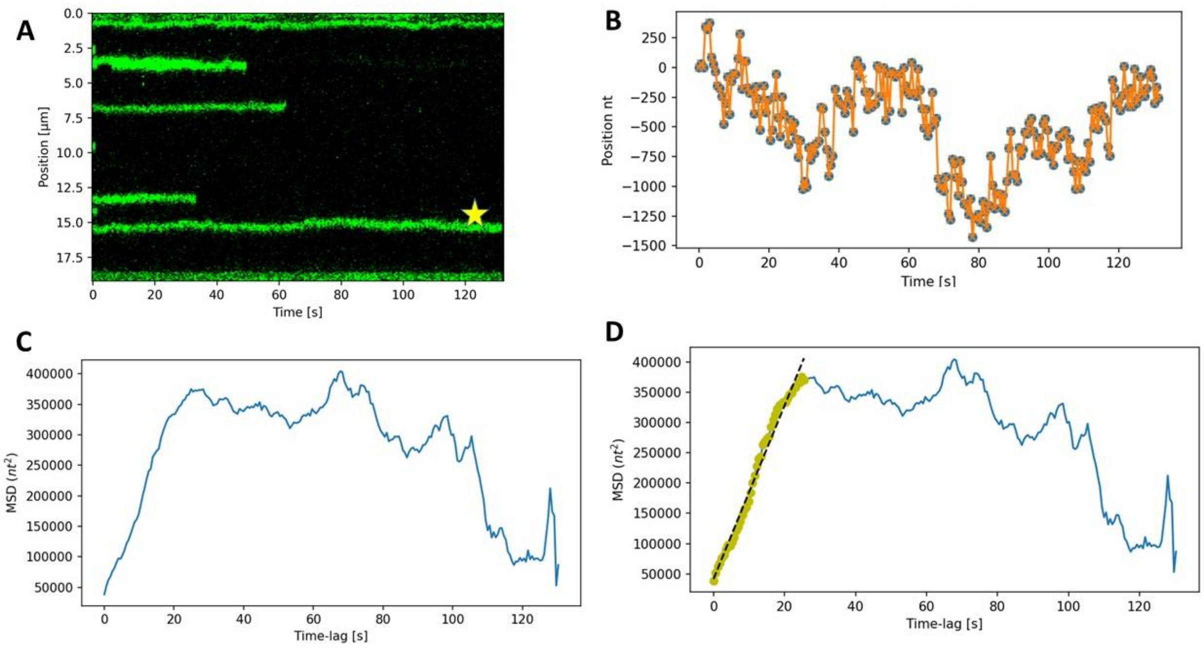


**Figure S1: Schematics of diffusion Calculation.** For individual diffusing proteins, we calculate the mean square displacement (MSD) and the linear portion of this is then fitted for 1D random diffusion with the equation MSD=2DT where the slope of the line would be diffusion coefficient(D). T in the equation is time. Given experiment is performed under 100 mM KCl [30 mM HEPES pH 7.8, 100 mM KCl, 6 % Glycerol, 5 mM $MgCl_2$] for RPA DBD D at 5 pN. The experiment was performed in a room temperature of 22°C. Yellow star indicates the protein trace used for the depicted analysis.

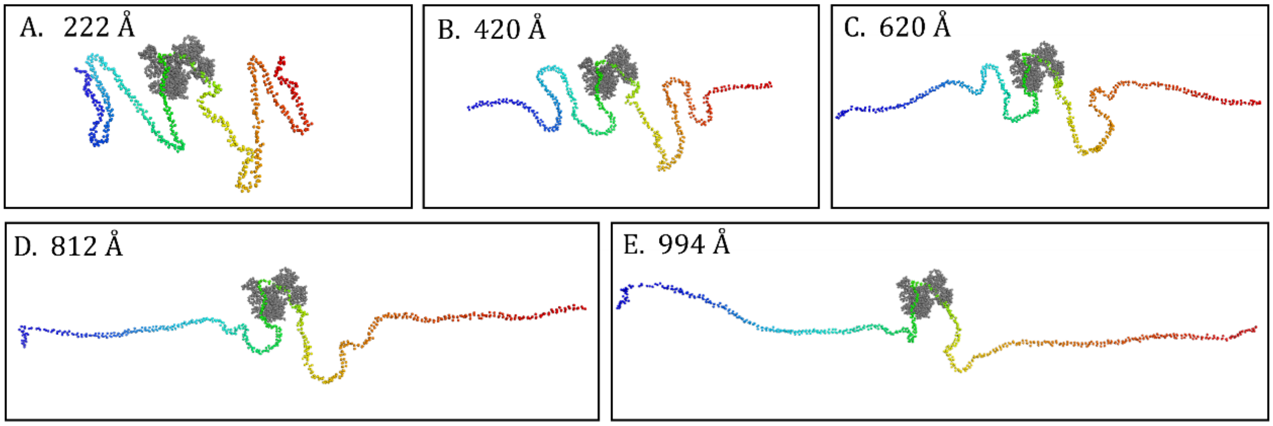


**Figure S2: Modelling force applied on the ssDNA ends.** Different initial conformations of the ssDNA with 235 nucleotides with end-to-end distances for these initial conformations are: 222 Å, 420 Å, 620 Å, 812 Å, and 994 Å. The RPA is shown in grey and is located at the center of the ssDNA.

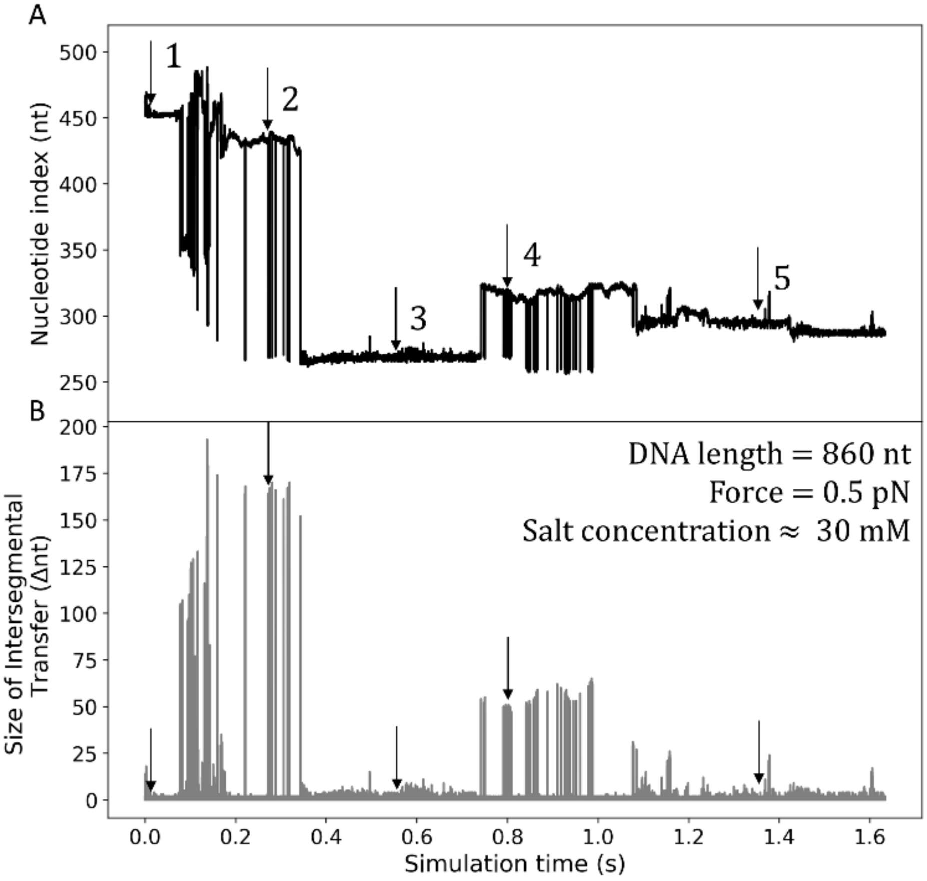


**Fig. S3**: **Diffusion of RPA on ssDNA through intersegmental transfer.** A. The variation of the nucleotide index (nt) closest to the centre-of-mass (COM) of the RPA protein is shown throughout a representative simulation trajectory for ssDNA of length of 860 nucleotides ssDNA with a force of 0.5 pN on the ssDNA (which corresponds to an end-to-end distance of 817 Å). The persistence length of the ssDNA is 32 that corresponds to salt concentration of 30 mM (which was modelled by force constant of 0.7 for the dihedral angles. B. Difference of the nucleotide index closest to the protein COM, *Dnt*, between two consecutive MD steps is shown with respect to the simulation time. The value of *Dnt* corresponds to loop of ssDNA bridged due to an intersegmental transfer event. C. Five representative snapshots of RPA-ssDNA binding sampled from the trajectory shown in (A). These five snapshots are designated as 1-5 and are indicated with arrows in A and B. On the right-hand side of each of the snapshots, the respective *Δnt* values are shown.

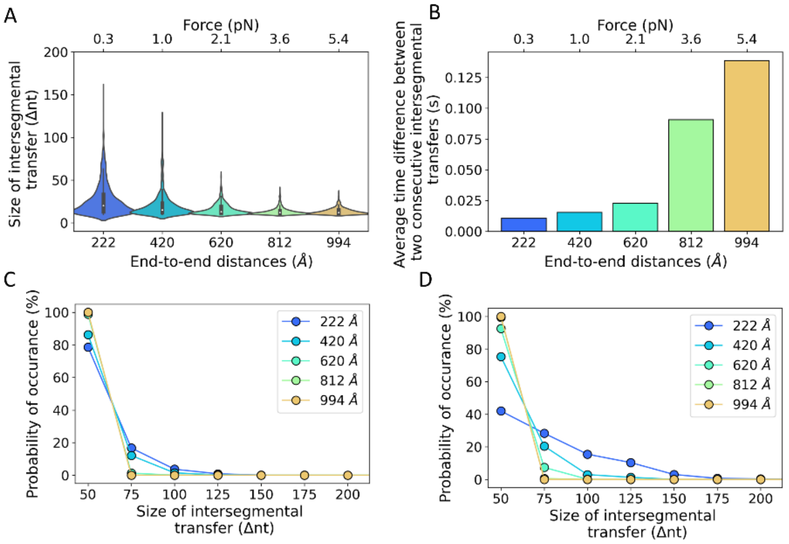


**Figure S4: Variation in the frequency of intersegmental transfer at different force and salt concentrations.** Characteristics of the intersegmental transfer events of RPA when diffusion along ssDNA under force applied on its ends. These simulations were performed at 1000 mM (panels A-C). A) Violin plots of the length of ssDNA, Δnt, skipped in intersegmental transfer of RPA when diffusing along ssDNA at five different forces applied on its two ends [the applied force (shown in the upper X axis), each of these forces corresponds to different end-to-end distance varying between 222 - 994 Å, (shown in the lower X axis)]. The simulations at different force applied on the two ssDNA ends are shown with different color. B) The kinetics of the intersegmental transfer is indicated by the average time difference between any two consecutive intersegmental transfer events (having length Δnt > 10) for the five systems. C) Distributions of the Δnt for systems simulated under different forces and high salt concentration (1000 mM). D) Distributions of the Δnt for systems simulated under different forces and at low salt concentration (30 mM).

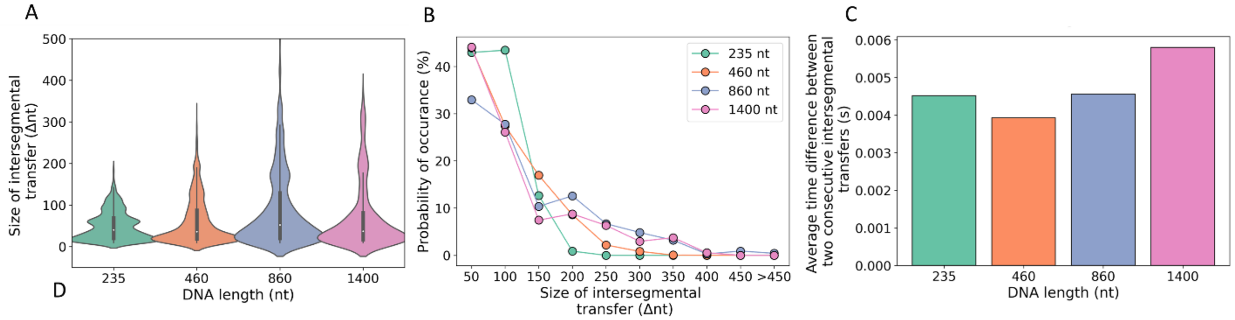


**Figure S5: Variation of RPA diffusion coefficient as a function of ssDNA length, and effect on intersegmental transfers.** A. Distribution of the length of intersegmental transfer at different DNA lengths. B. The occurrence of different sizes of intersegmental transfer during a simulation trajectory as a function of DNA length. C. The average time difference between any two consecutive intersegmental transfers (having length Δnt > 10) for four different DNA lengths.

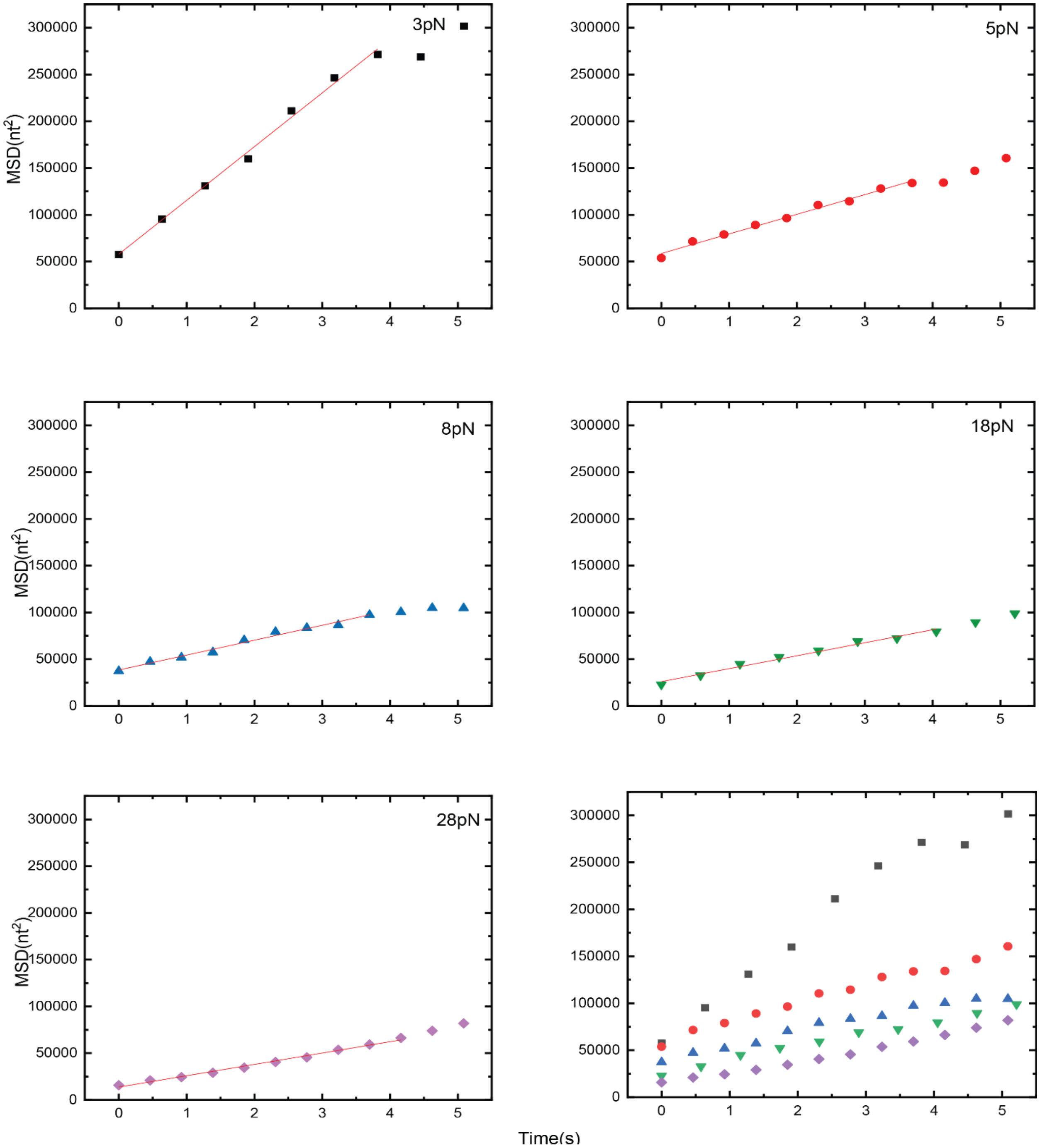


**Figure S6: The time course of mean squared displacement (MSD) at various forces.** Forces are indicated in the plot. The linear section is fitted to a linear equation, and the obtained slope divided by two represents the 1D diffusion coefficient. The fitted lines were drawn in respective colors in dash lines.

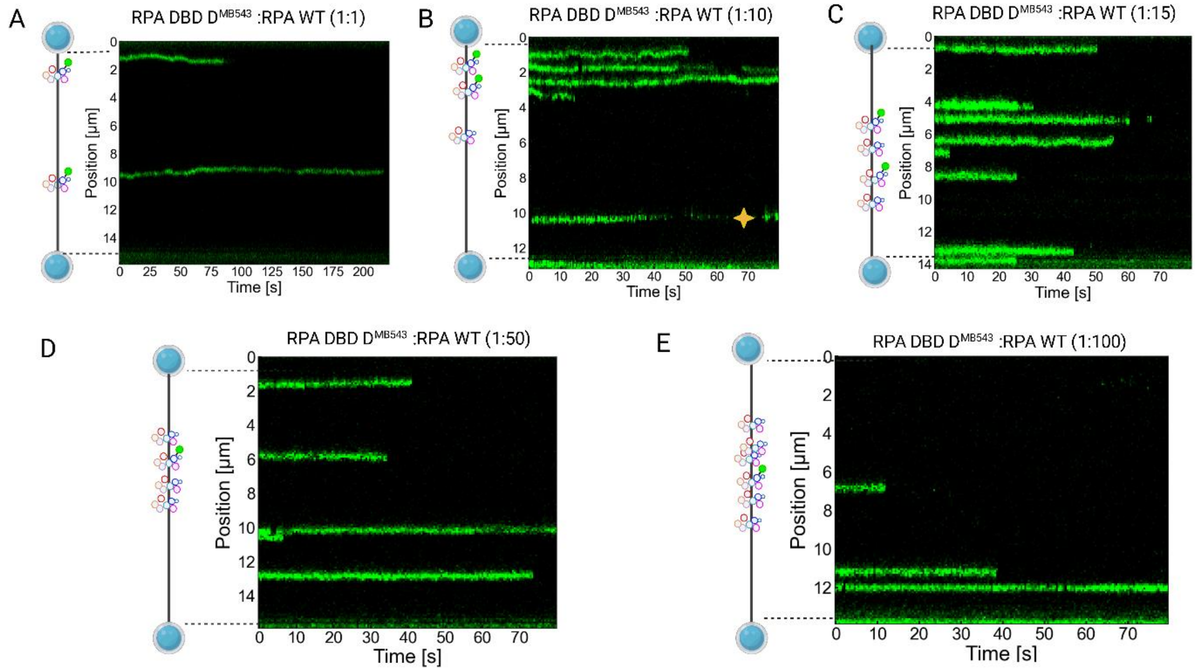


**Figure S7: Crowding effect as depicted in kymograph.** A yellow star indicates the aggregates which are excluded from data analysis. All experiments were performed under 5pN tension on DNA and 100 mM KCl condition. Green circle for the RPA cartoon represents the label while missing green circle represents unlabeled RPA.RPA DBD D concentration was 10 pM and RPA WT concentration was adjusted as per the ratio. Labeled and unlabeled samples were pre-mixed before the experiment and the waiting time for the binding of proteins was 30 s unless otherwise mentioned.

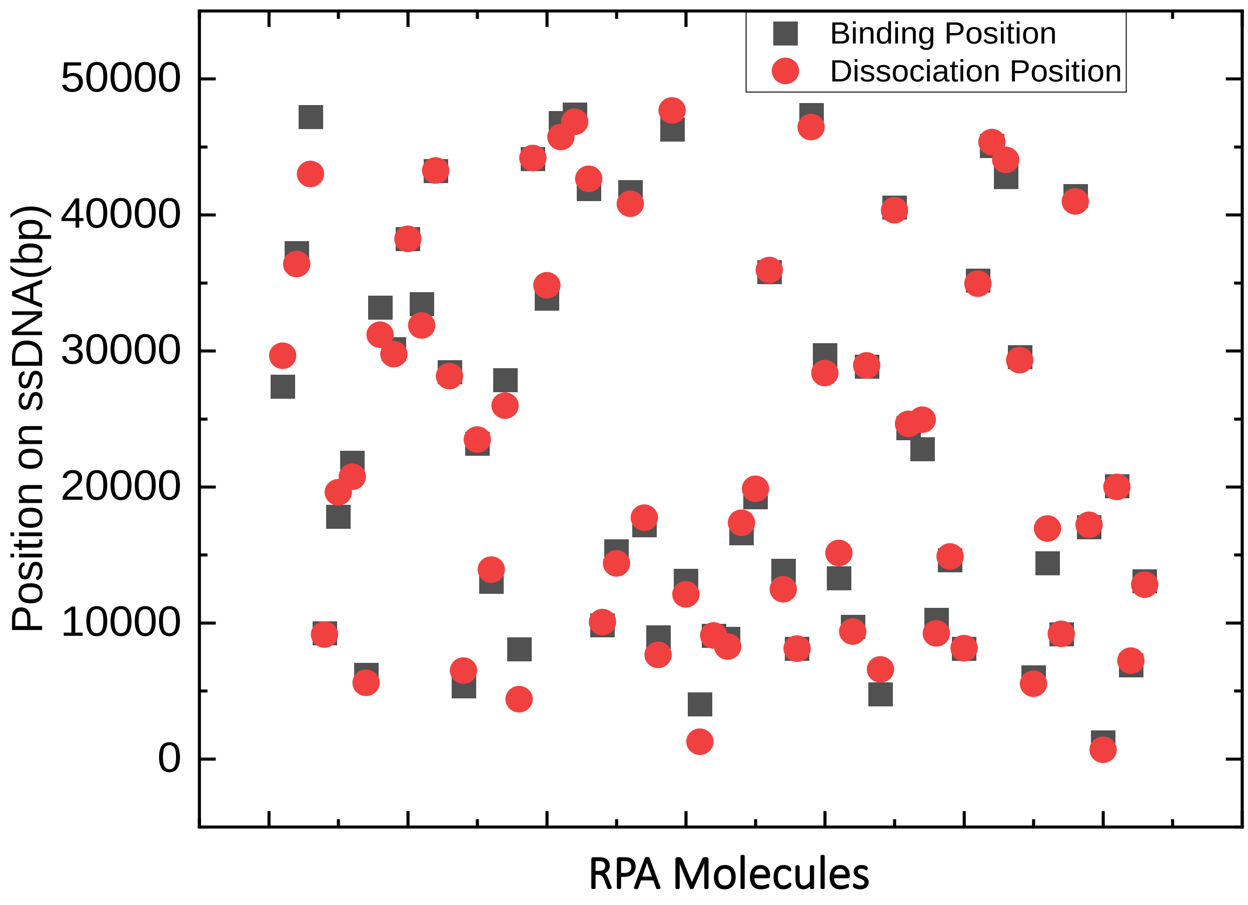


**Figure S8: Binding and dissociation on genomic position shows no specific binding or dissociation.** Initial binding position of RPA on lambda ssDNA and dissociation position on lambda DNA for individual traces of RPA DBD at 100 mM KCl were looked under 5 pN stress. Black square represents binding position while red circle denotes dissociation position. We see that there is no specific accumulation at certain position indicating no sequence specificity of RPA.

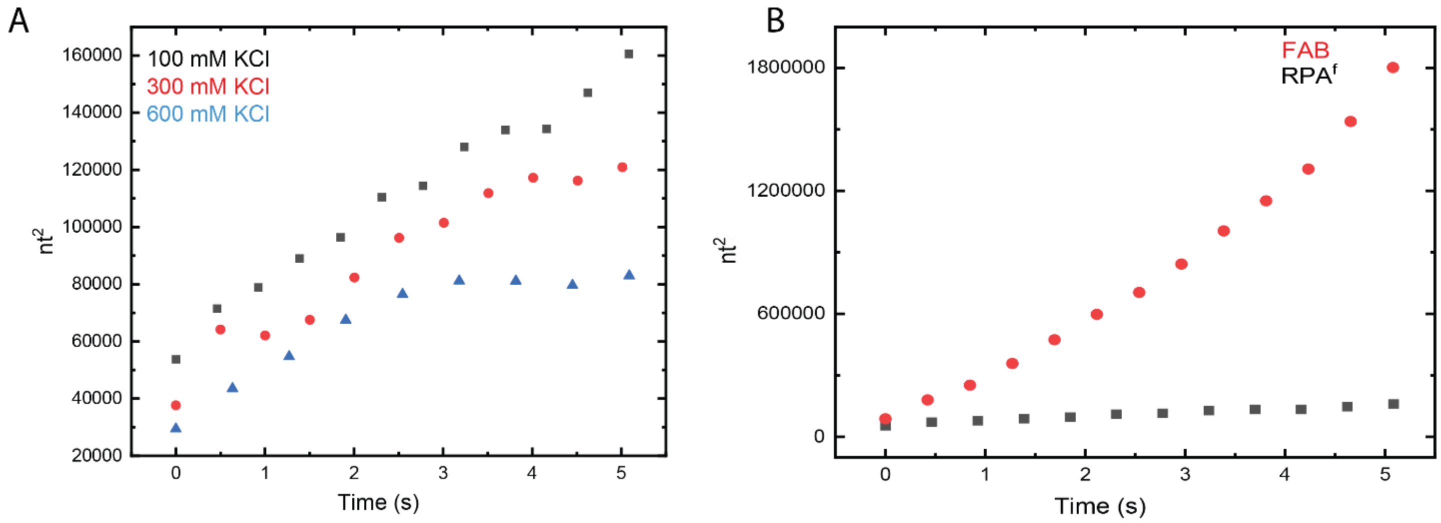

**Figure S9: MSD vs Time for RPA WT at different salt concentration.** MSD vs Time for RPA full length FAB at 5pN. A) MSD vs Time at 5 pN for different salt concentration as indicated by different colors. A linear line was fitted to calculate the diffusion coefficient. B) MSD vs Time for the full-length RPA vs FAB. A linear line was fitted to calculate the diffusion coefficient.

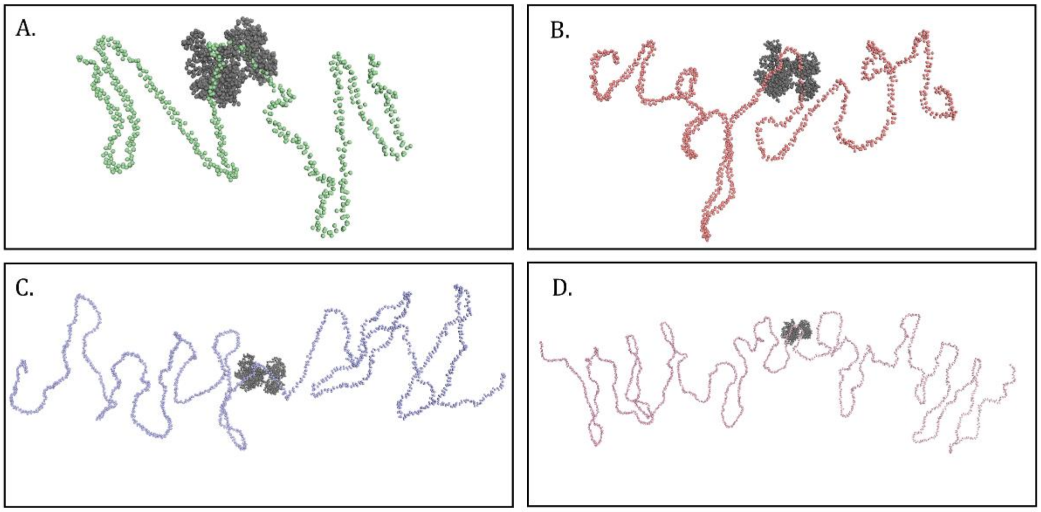


**Figure S10: Different DNA lengths.** Initial conformations of the RPA interacting with different lengths of ssDNA: A. 235 nucleotides, B. 460 nucleotides, C. 860 nucleotides, D. 1400 nucleotides.

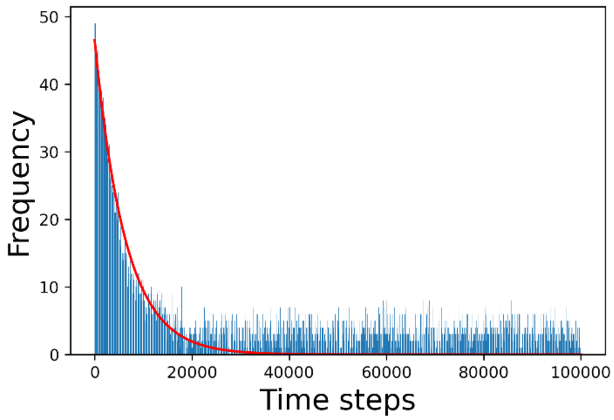


**Figure S11: Frequency of unfolded state lifetime:** Frequency of unfolded state as a function of MD time steps

To convert the MD simulation time steps to real time unit, a short fragment of DNA (20 nucleotides) has been simulated in space where we have applied contact interaction between five base beads from both ends. After certain time steps the contacts are formed and the DNA fragment goes from an unfolded conformation to a folded conformation. We define a state as folded state if at least three contacts are formed among these five pairs of beads. With this identical condition, 50 simulations were run. Then, we calculated the number of folded and unfolded state at every simulation time steps from these simulations. Finally, we plotted frequency (or number) of unfolded states as a function of simulation time steps. We obtained the unfolded state lifetime from this plot as 6290.20 simulation time steps which corresponds to the experimental value of $2 \times 10^{-3}$ s. This plot is similar to that obtained from the works of Kim et al (80). Thus, we matched this experimental unfolded state lifetime with that obtained from the simulations (80). Since we used a shorter fragment than the one used experimentally, we extrapolated the relevant lifetime given the experimental relationship between the lifetime and length of the ssDNA (80). Using this relationship, the relevant lifetime in seconds for 20 nucleotides ssDNA was obtained. From this, we obtained the equivalent time for 1 MD time steps which is equal to $3.267 \times 10^{-8}$ s.

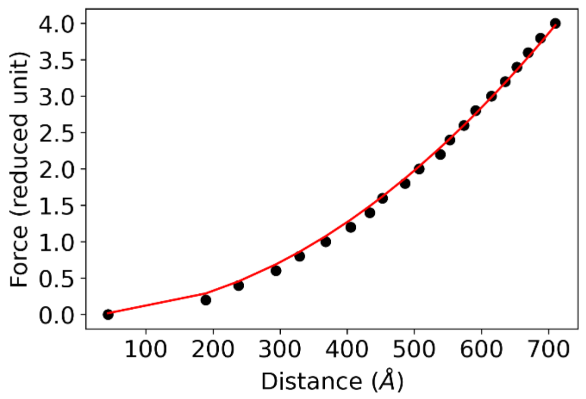


**Figure S12: Force extension curve for current model:**

To convert the modeled end-to-end distances to corresponding force values, we took an ssDNA polymer of length 235 nucleotides and applied various values of force at its two ends ranging from 0.2 to 4.0 reduced units. We ran short simulations of $1 \times 10^6$ timesteps for each case and calculated the end-to-end distances at the end of the simulations. Now, we plot the force values as function of the end-to-end distances. This plot was fitted to FJC model given by,

$$x = L_0[\coth(\frac{2FL_p}{k_BT}) - \frac{k_BT}{2FL_p}](1 + \frac{F}{K_0})$$

From the fitting, we obtained the values of $L_0$ and $K_0$, where $K_0$ is the elastic modulus of the polymer, $L_0$ is the contour length of the polymer and $L_p$, $k_B$ and T being the is the persistence length, Boltzmann's constant, the simulation temperature respectively, while F and x are the force and end-to-end distances. After obtaining this $L_0$ value we normalized it with the number of nucleotides in the system. And with the normalized $L_0$ value and $K_0$, we obtained the force value for different systems. For different salt concentrations we utilized different $L_p$ values.

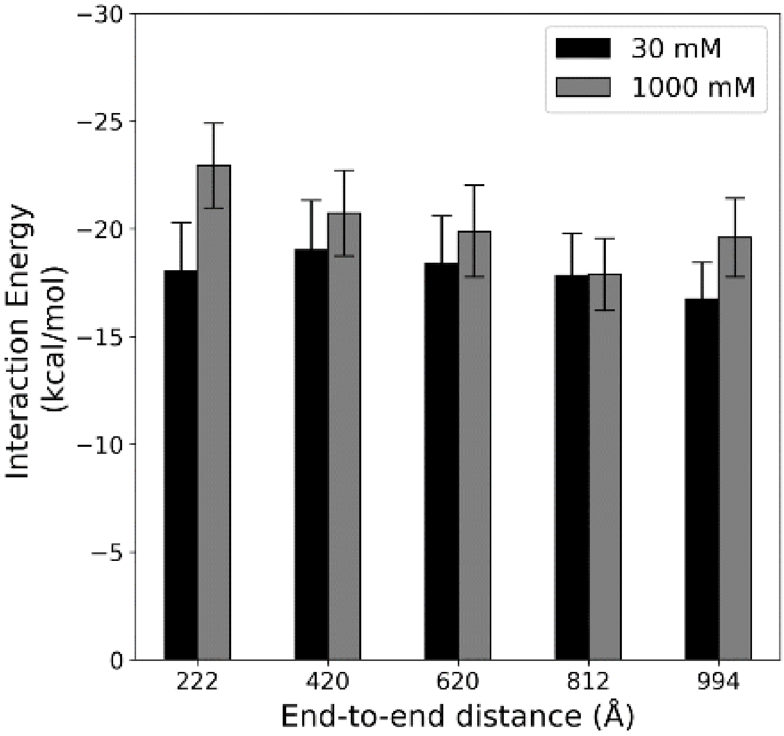


**Figure S13: Variation of diffusion coefficient at different force and salt concentrations.** A bar diagram representing RPA-ssDNA interaction energies for five different forces applied on the ssDNA ends (i.e., different end-to-end distances) at two salt concentrations (black and grey).
